# DrugRepo: A novel approach to repurpose a huge collection of compounds based on chemical and genomic features

**DOI:** 10.1101/2022.04.21.488995

**Authors:** Yinyin Wang, Jehad Aldahdooh, Yingying Hu, Hongbin Yang, Markus Vähä-Koskela, Jing Tang, Ziaurrehman Tanoli

## Abstract

The drug development process consumes 9-12 years and approximately one billion US dollars in terms of costs. Due to high finances and time costs required by the traditional drug discovery paradigm, repurposing the old drugs to treat cancer and rare diseases is becoming popular. Computational approaches are mainly data-driven and involve a systematic analysis of different data types leading to the formulation of repurposing hypotheses. This study presents a novel scoring algorithm based on chemical and genomic data types to repurpose vast collection of compounds for 674 cancer types and other diseases. The data types used to design the scoring algorithm are chemical structures, drug-target interactions (DTI), pathways, and disease-gene associations. The repurpose scoring algorithm is strengthened by integrating the most comprehensive manually curated datasets for each data type. More than 100 of our repurposed compounds can be matched with ongoing studies at clinical trials (https://clinicaltrials.gov/). Our analysis is supported by a web tool available at: http://drugrepo.org/.

## 1. INTRODUCTION

The average cost of developing a new drug is billions of dollars, and it takes about 9–12 years to bring a new drug to the market ^[1]^. Finding new uses for approved drugs has become a primary alternative strategy for the pharmaceutical industry. This practice, usually referred to as drug repositioning or drug repurposing, is highly attractive because of its potential to speed up the process of drug development, reduce costs, and provide treatments for unmet medical needs ^[2]^. In this regard, compounds that have passed through phases I or II in the drug discovery pipeline but never made it to the market due to efficacy issues carry great potential for drug repositioning. Traditionally, drug repurposing success stories have mainly resulted from largely opportunistic and serendipitous findings ^[3]^. For example, sildenafil citrate was originally developed as an antihypertensive drug but later repurposed by Pfizer and marketed as Viagra to treat erectile dysfunction based on retrospective clinical experience, leading to massive worldwide sales. Other examples of such drug repositioning include cancer drugs: crizotinib, sorafenib, azacitidine and decitabine, all of which failed to reach the markets in their initial indications yet now are essential tools in the treatment of various types of cancers ^[4]^.

Over the recent years, various computational resources are developed to support systematic drug repurposing. Popular information sources for *in-silico* drug repurposing include, for instance, electronic health records, genome-wide association analyses or gene expression response profiles, pathway mappings, compound structures, target binding assays and other phenotypic profiling data ^[3]^. Several systematic review articles on the use of computational repurposing approaches are available that cover machine learning (ML) algorithms ^[5][6][7]^. Several databases directly support *in-silico* drug repurposing, including Drug Repurposing Hub ^[8]^, repoDB ^[9]^ and RepurposeDB ^[10]^. On the other hand, hundreds of databases can indirectly support drug repurposing ^[7][11]^. However, these databases provide experimentally tested indications only for a limited number of investigational or approved compounds and ignore the massive number of preclinical compounds that could be potential candidates for drug repurposing. Drug target profiles for approximately two million such preclinical compounds are available at ChEMBL^[12]^ and other databases.

Drug-target interactions (DTI), meaning the target molecules each compound binds to and the relative binding strength and impact on cellular functions, lie at the heart of drug discovery and repositioning. Several artificial intelligence (AI) methods for drug repurposing are based on DTIs as well as chemical structural similarities ^[13][14][15]^. However, these methods are applied only to a selected set of compounds resulting in limited prediction outcomes ^[13]^. Computational approaches are primarily data-driven and involve a systematic analysis of several components (or data types) before suggesting a repurposed indication. These components may include chemical structures, adverse event profiles, compound-target interactions, pathways, disease-gene associations, genomic, proteomic, and transcriptomic information. The drug repurposing methods can be developed based on the individual or combination of these components.

In this study, we propose DrugRepo (http://drugrepo.org/); a novel scoring algorithm that can effectively repurpose hundreds of thousands of compounds based on three components, 1) overlapping compound-targets score (OCTS), 2) structure similarity score based on Tanimoto coefficient (TC), and 3) compound-disease score (CDS). The DrugRepo score is computed between the approved drug (for a particular disease) and candidate compound and is the average of the three component scores. Approved indications for **674** diseases and **1**,**092** compounds are collected from https://clinicaltrials.gov/. To explore the translational impact of DrugRepo, we cross-referenced candidate compounds with completed clinical trials at https://clinicaltrials.gov/. We observed that 186 compounds are explored in different clinical studies across nine cancer types. We also compared our candidate compounds with the predicted compound disease relationships at the Comparative Toxicogenomic Database (CTD) ^[16]^ and found a statistically significant overlap. These promising findings demonstrate the versatility of DrugRepo. Our new tool provides a quick and effective scoring method for drug repurposing.

## 2. MATERIALS

Several types of data are integrated into this analysis, e.g., approved drug indications, compound-target profiles, disease-gene associations, and protein-protein interaction (PPI) networks. These datasets are consequently explained in the following subsections.

### 2.1. Approved drug indications

Approved drug indications are extracted from the clinical trials database (https://clinicaltrials.gov/), as it is the most up to date repository for drug indications and clinical phases for the compounds. However, the data provided by clinical trials is not well structured and doesn’t provide standard naming conventions or identifiers for the compounds and diseases. We, therefore, utilized a semi-automated approach to extract drug-disease indications, assigned UML-CUI and standard InChIKey identifiers for drugs and diseases, respectively. The standard InChIKey mapping is performed using the PubChem python client (https://pubchempy.readthedocs.io/en/latest/), whereas UML-CUIs are assigned to the diseases using disease annotations provided by DisGeNET ^[17]^. Finally, we extracted data for 674 diseases, 1,092 drugs, and 3,868 approved drug indications, as shown in Supplementary file 1.

### 2.2. Compound-target profiles

Non-overlapping compound-target profiles are extracted from the five most comprehensive (and manually curated) databases namely, ChEMBL ^[18]^, BindingDB ^[19]^, GtopDB ^[20]^, DrugBank ^[21]^ and DGiDB ^[22]^. The compound identifiers in these databases are mapped into standard InChIKeys and SMILES using UniChem ^[23]^ and PubChem ^[24]^, respectively, whereas identifiers for target proteins are mapped to UniProt identifiers ^[25]^. The combined non-overlapping compounds from the five databases exceed **2M** with approximately **15**,**000** targets. These comprehensive datasets are extracted using application programmable interfaces (APIs), standalone text files, and SQL dumps. The first three databases, ChEMBL, BindingDB and GtopDB, provide quantitative bioactivity data, such as measurements in terms of IC_50_, K_d_, and K_i_, whereas DrugBank and DGiDB contain unary but experimentally verified compound-target interactions. In addition to active or potent compound-target profiles in ChEMBL, BindingDB and GtopDB, there exists a big proportion of in-active compound-target profiles (concentration > 10,000 nM). These in-active compound-target profiles could jeopardize the analysis in the proposed research. Therefore, in this analysis, we considered only potent compound-target profiles (concentration is <=1000 nM) ^[26]^. Hence, we left with **788**,**078** compounds and **8**,**754** protein targets. Potent target profiles for these ∼0.8M compounds are already integrated and publicly available in MICHA (https://micha-protocol.org/) ^[27]^.

### 2.3. Disease-gene associations and PPI networks

To support the large-scale drug repurposing, we integrated manually curated disease-gene associations from DisGeNET ^[17]^. There are 9,703 genes, 11,181 diseases and 84,038 associations. These curated disease-gene associations are provided in Supplementary file 2.

Protein-protein interactions (PPI) networks were extracted from a manually curated human interactome, including 16,677 proteins and 243,603 PPIs ^[28]^.

## 3. METHODS

There are 788,078 compounds for which there exists at least one potent target (concentration is ≤1000 nM) in any of the five DTI databases. We call these agents candidate compounds to be repurposed. For each candidate compound, the DrugRepo score is calculated as the average of three component scores, OCTS, TC, and CDS, which are derived by comparing each candidate compound to 1,092 approved drugs (**Figure 1**). Because the number of calculated scores (788,078 × 1092) is too big for the web portal to handle smoothly, we considered only those cases where the structural similarity between the approved drug and candidate compound is ≥ 0.2. This way, we were left with 2,207,367 scores.

**Figure 1:**
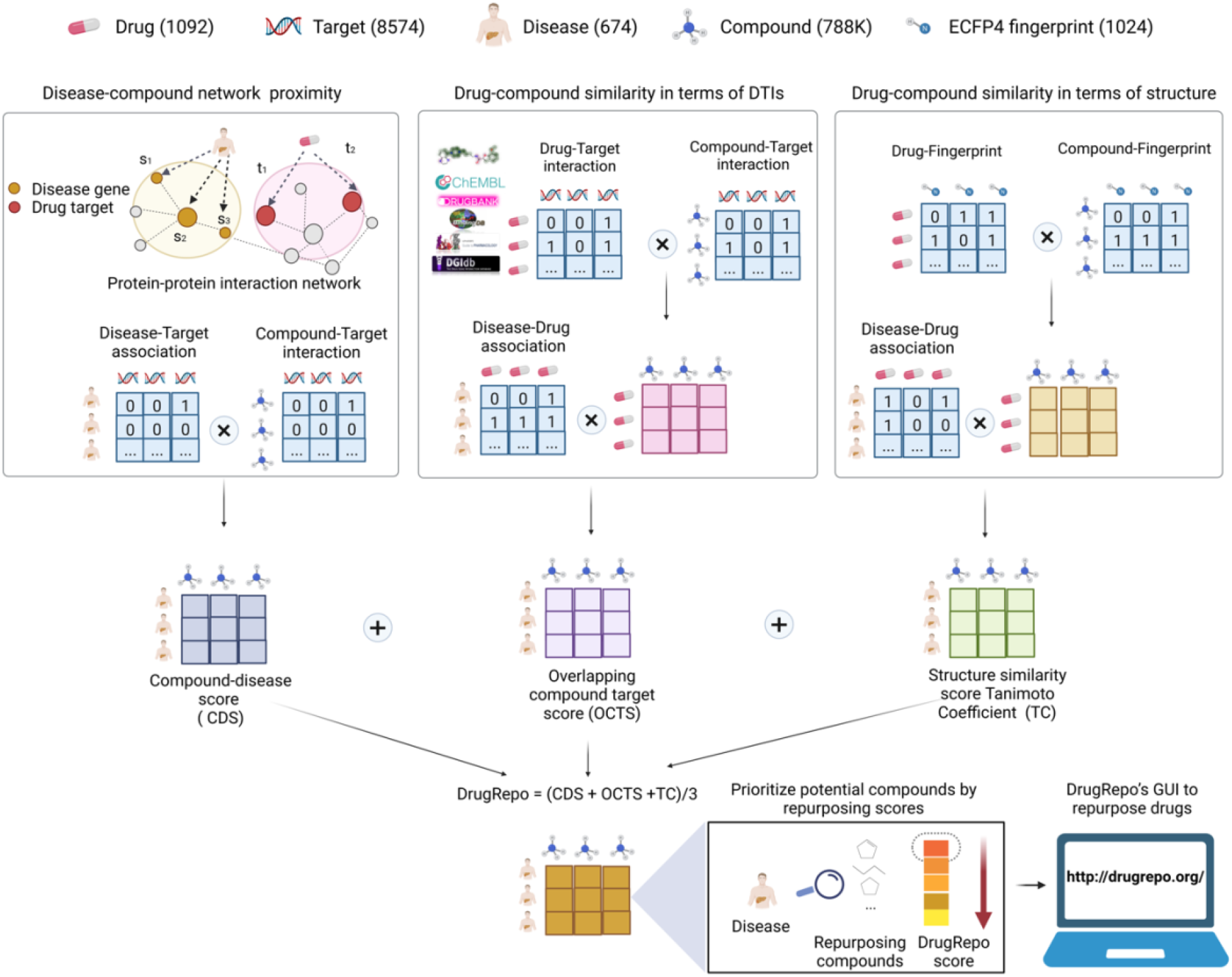
The schematic figure for drug repurposing in DrugRepo. The DrugRepo pipeline starts with the user selecting a particular disease. There are 0.8M candidate compounds in DrugRepo that can be repurposed for 674 diseases. At first, the pipeline finds approved drug(s) for the selected disease and searches for structurally similar compounds. In this step, the Tanimoto coefficient (TC) describes the structural similarity between molecular fingerprints (ECFP4) of approved and candidate compounds. A threshold is used to favor similar molecular structures. The second step is to compute DTI profiles for candidate compounds and approved drugs. The OCTS is the score based on overlapping DTIs between approved and candidate compounds. In case of multiple approved drugs for a disease, we took average of OCTS and TC scores. The third step is to compute the compound-disease score (CDS). The CDS is the average of the minimum distances in the PPI networks between target molecules and molecules associated with the selected disease. The average distance is normalized to 0-1. Finally, the DrugRepo score is calculated as the average of the three component scores. The higher the DrugRepo score between the approved drug and the candidate compound, the higher the possibility of repurposing the compound for a particular disease. Finally, we developed the DrugRepo’s GUI to provide a user-friendly service for repurposing drugs with our pipeline. The OCTS between approved and candidate compounds are computed using equation 1. The OCTS ranges from 0 to 1 and represents the proportion of targets shared between an approved drug and the candidate compounds. Candidate compounds sharing more targets with the approved drugs will have higher OCTS values.

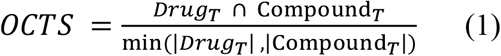

Where *Drug*_*T*_ *and* Compound_*T*_ are the sets of potent targets for a pair of approved drugs and candidate compounds, respectively. Similarly, |*Drug*_*T*_|) *and* |Compound_*T*_|) are the total number of targets associated with approved drug and the candidate compound respectively.

Compound-target profiles are extracted from five databases. The number of overlapping compounds and targets in these five databases are shown in **Figure 2A** and **Figure 2B**, respectively. As evident from **Figure 2A**, ChEMBL features the most comprehensive collection of compounds, whereas DrugBank covers more significant number of targets (**Figure 2B**). There are 483 compounds and 776 targets that are common in all five databases.

**Figure 2:**
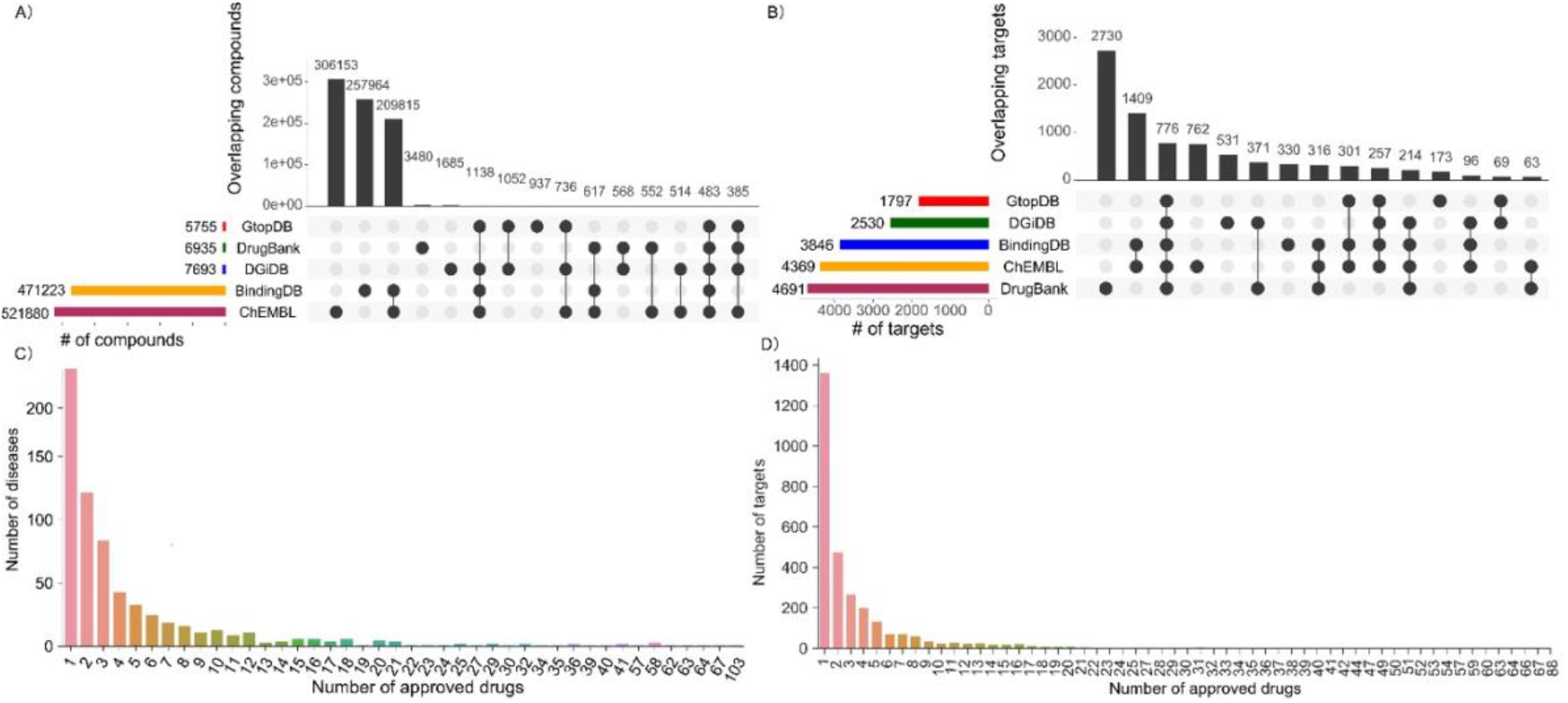
(A) Number of overlapping compounds across five databases. (B) Number of overlapping targets between five databases. (C) Distribution of diseases across 1092 approved drugs. (D)Target coverage across 1092 approved drugs.

Drug repurposing is challenging because of shortcomings in data coverage. The diseases associated with significant number of approved drugs may have better chances of correctly repurposing the compounds as the number of candidate compounds will also be larger. However, only very few diseases are associated bigger number of approved drugs. HIV is associated with the highest number of approved drugs (n = 103), but as shown in **Figure 2C**, more than 70% of diseases have less than five approved drugs. On the other hand, the lack of drug-target interactions is also a hurdle as it limits matching of compounds by the putative mechanism of action. Indeed, most approved drugs have less than 30 targets (**Figure 2D**). To compensate for the shortage of approved drugs and drug-target-interactions, we incorporated two additional components in the DrugRepo pipeline: the Tanimoto coefficient (TC), which is a structural similarity score, and the compound-disease score (CDS), which ranks new compounds based on how closely their target spaces match with the target proteins that are associated with the disease.

The Tanimoto coefficient (TC) is measures structural similarities between molecular ECFP4 fingerprints of approved and candidate compound for a particular disease. The fingerprints are the bit strings denoting the presence or the absence of chemical substructures and are calculated using RDKit package [29].

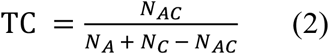

Where *N*_*A*_ *and N*_*C*_ are the number of sub-structures present in the approved drug and candidate compound, respectively, and *N*_*AC*_ are number of common sub-structures found in both approved drug and the candidate compound. The value of TC is between range 0-1 and constitutes the second component of DrugRepo.

The compound-disease score (CDS) is measured by averaging the minimum distances in PPI networks between potent targets of the candidate compound and genes associated with the disease, as shown in equation 3. Curated disease-gene associations were extracted from DisGeNET for the 674 diseases used in this analysis. There are 22,399 non-zero CDS values.

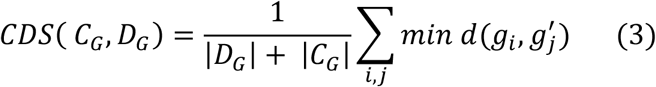

Where *C*_*G*_ = (*g*_1_,*g*_2_,…) is the set of gene targets for candidate compounds and 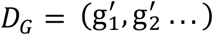 are the genes associated with a particular disease (acquired from DisGeNET). The average of minimum distances between *C*_*G*_ *and D*_*G*_ are computed in PPI networks. The average distance is further normalized to 0-1 using min-max normalization. Finally, the DrugRepo score is the mean of the three compound scores and ranges from 0 to 1. The higher the DrugRepo score between an approved drug (for a particular disease) and a candidate compound, the greater the repurposing potential of the candidate compound for that disease.

## 4. RESULTS AND DISCUSSIONS

To explore the translational impact of DrugRepo, we evaluated our repurposed compounds using two methods, i.e. 1) cross-referenced thousands of the repurposed compounds using disease-compound associations in CTD ^[16]^, 2) matched 186 compounds across nine cancer types for which either Phase I or Phase II trials have been completed or Phase III trials is ongoing.

### 4.1. Matching repurposed compounds with disease-compound associations in CTD

The Comparative Toxicogenomic Database (CTD) contains manually curated and inferred compound-disease relationships ^[16]^. CTD associates thousands of compounds with diseases based on drug-target and disease-gene relationships. Our scoring method and the datasets are different from CTD, but the output types in DrugRepo and CTD are the same. We, therefore, assessed the accuracy of DrugRepo by comparing the repurposed compounds with compound-disease relationships in CTD. We downloaded disease-compound relationships from CTD at: http://ctdbase.org/downloads/;jsessionid=5DA98FA00707F8A41A59335C1CC36C47#cd. In CTD, compounds are represented by CAS identifiers or compound names; and diseases are represented using Mesh ids. To make the comparison possible, we mapped compound names from the CTD dataset into standard InChIKeys using PubChempy. Diseases are mapped from MESH ids into UML-CUI. There are 1,048,548 compound-disease associations in CTD, including 3,941 compounds and 6,119 diseases. Many of these compounds and diseases are unable to map into standard InChIKeys and UML-CUI identifiers. We, therefore, skipped those cases and left with only 168,471 compound-disease associations (605 diseases and 2,598 compounds), as shown in Supplementary file 3. These associations are used to match repurposing candidate compounds by DrugRepo. The significance of matching results for overlapping compounds (for a particular disease) between DrugRepo and CTD is computed using the following equations:

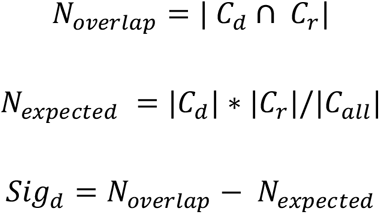

Where *C*_*all*_ is the set of all candidate compounds (∼0.8M) in DrugRepo, *C*_*d*_ is set of compounds associated to disease ‘d’ in CTD, *C*_*r*_ is the set of repurposed compounds by DrugRepo for the same disease ‘d’, *N*_*overrlap*_ is the number of overlapping compounds between CTD and DrugRepo, *N*_*expected*_ is the expected number of compounds overlapping with *C*_*d*_ if we randomly choose the same number of compounds (|*C*_*r*_|) from *C*_*all*_, and *Sig*_*d*_ is the significance of matched compounds. If the significance is greater than 0, it means our repurposing is not random.

We have ∼0.8M candidate compounds and 1,092 approved drugs. This could result in a vast matrix (∼800M). Therefore, we considered only those candidate compounds that at least have 50% structural similarity with any of the approved drugs (TC >= 0.5). If *N*_*overrlap*_ for a particular disease is greater than *N*_*expected*_, *Sig*_*d*_ will be positive, meaning that repurposed compounds by DrugRepo are matched with CTD. Diseases with the significance scores > 0 are shown in **Figure 3A** (*N*_*expected*_ is shown with blue and *N*_*overrlap*_ with red bars). As shown in **Figure 3A**, the blue bar represents the expected number of compounds, and the red bar indicates the actual number of matched compounds. Most blue bars have significance scores less than 1, suggesting that if chosen randomly, there should be less than one candidate compound that can match for most of the diseases. We matched more than one correctly repurposed compound for 118 diseases (**Figure 3A**). In other words, DrugRepo shows good repurposing ability in 118 out of the 605 diseases in CTD, suggesting that our novel pipeline effectively repurposes compounds for several diseases. For example, DrugRepo returns 4,921 candidate compounds and 401 compounds by CTD for myocardial infarction. If the selection of the 4,921 compounds (from the 788,078 total candidate compounds) had been random, we would have expected an average of 2.51 matched compounds. However, we matched 15 compounds with CTD (6 folds bigger than the expected number), suggesting that the DrugRepo pipeline extracts biologically relevant candidate compounds.

**Figure 3:**
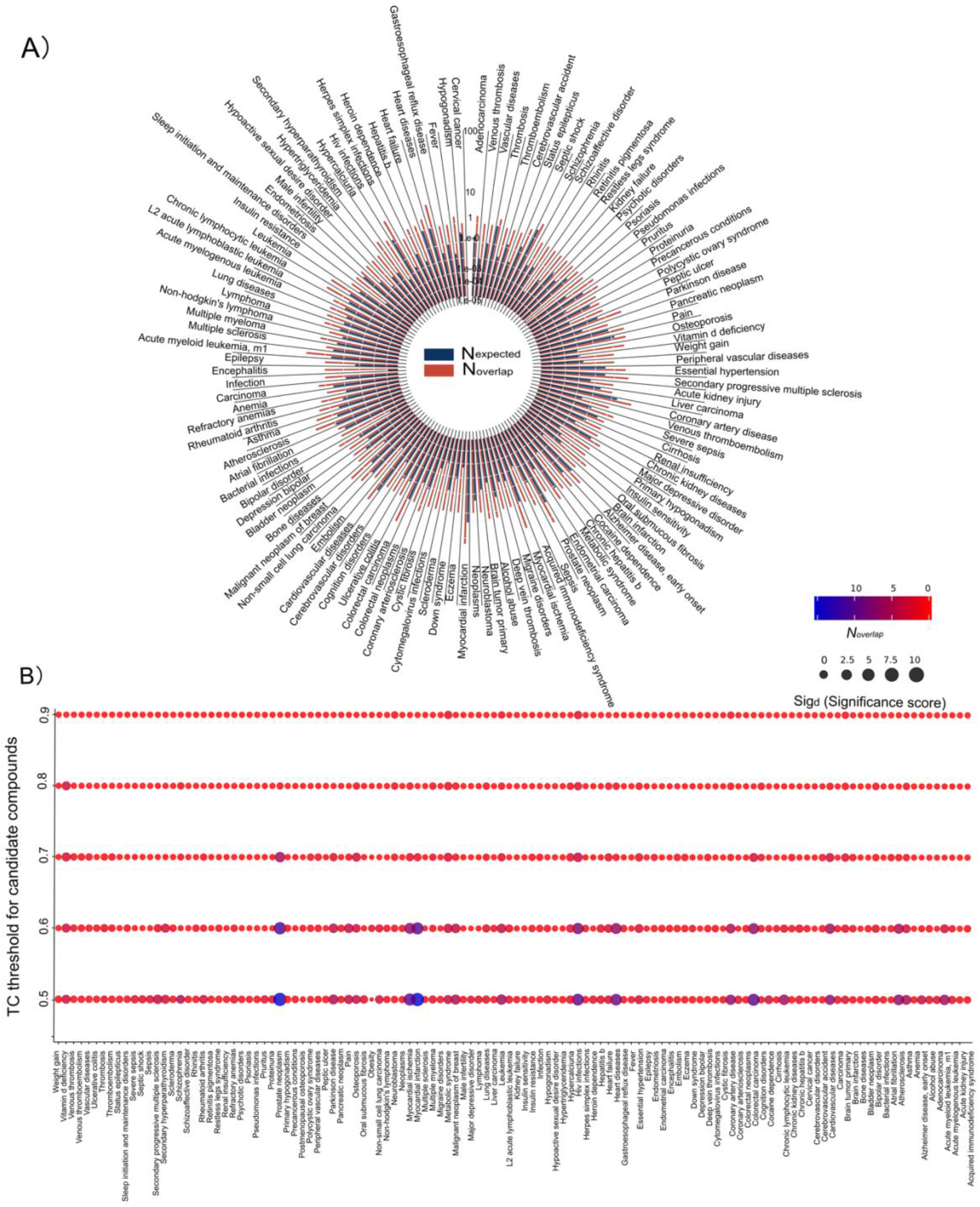
(A) The circular bar shows drug repurposing results matched with CTD. All the matched diseases were distributed as pie in the circle. The blue bars represent (N_expected) the expected number of compounds, if chosen randomly while the red bar represents the actual overlap (Noverlap) between compounds in CTD and DrugRepo. (B) The Significance scores at different TC thresholds (with y-axis as thresholds and x-axis as a disease whose true positive is not 0). The significance score is represented by dot size, and the colour from red to blue represents number of overlapping compounds.

To investigate the effect of structural similarity on drug repurposing, we evaluated the matched repurposed compounds on five different thresholds (TC: 0.5, 0.6, 0.7, 0.8, 0.9). As shown in **Figure 3B**, the number of matched repurposed compounds tends to decrease with strict TC filtration on repurposed compounds, as expected. However, the significance scores are also reduced, especially after TC ≥ 0.9, suggesting that high structure similarity is not a determining factor for drug repurposing. Many of the matched repurposed compounds are located at **TC >= 0.5**. On the other hand, the number of matched compounds and significance scores is relatively stable between 0.6 ≤ TC ≤ 0.7. Therefore, TC values between 0.6 to 0.7 might be optimal for drug repurposing.

We also analysed whether diseases associated with a more significant number of approved drugs can affect the DrugRepo scoring. As shown in Figure 4A, if a specific disease is associated with a considerable number of approved drugs, then more repurposed compounds can be matched (correlation = 0.7). Similarly, the number of matched repurposed compounds (N_overlap_) is also closely associated with the significance score (**Figure 4B**). Conversely, DrugRepo performance remains poor for the diseases associated with fewer approved drugs.

**Figure 4:**
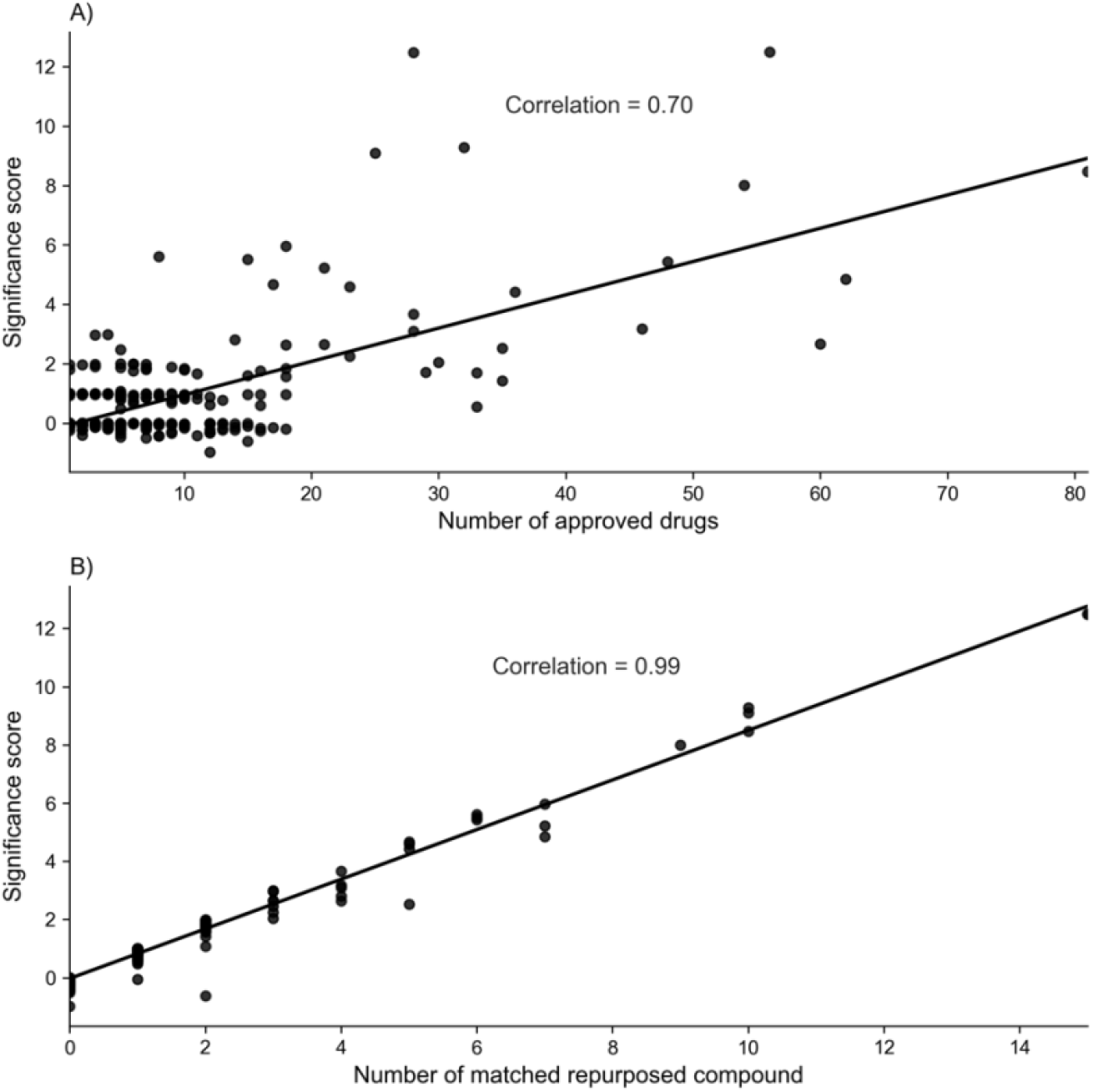
The impact of significance score on number of: (A) approved drugs associated with a disease, (B) matched repurposed compounds.

### 4.2. Matching DrugRepo candidate compounds with drugs in clinical trials

To establish if the DrugRepo pipeline enriches for compounds already tested in patients, we chose nine commonly studied cancer types to match repurposed compounds in DrugRepo with completed Phase I or Phase II or ongoing Phase III clinical trials. First, using the clinical trial API (https://clinicaltrials.gov/api/), we obtained the names of all compounds matching these criteria. Using PubChem’s API, we then tried to map the compound names with standard InChIKeys. However, the naming convention for compounds is not yet standardized, and several compounds were not mapped and therefore omitted from the subsequent steps.

**Figures 5A** and **5B** showed that we found 186 and 51 candidate compounds using structural similarity thresholds of TC >= 0.2 and TC >= 0.5, respectively. The names and identifiers of these matched compounds and other statistics are provided in Supplementary file 4.

**Figure 5:**
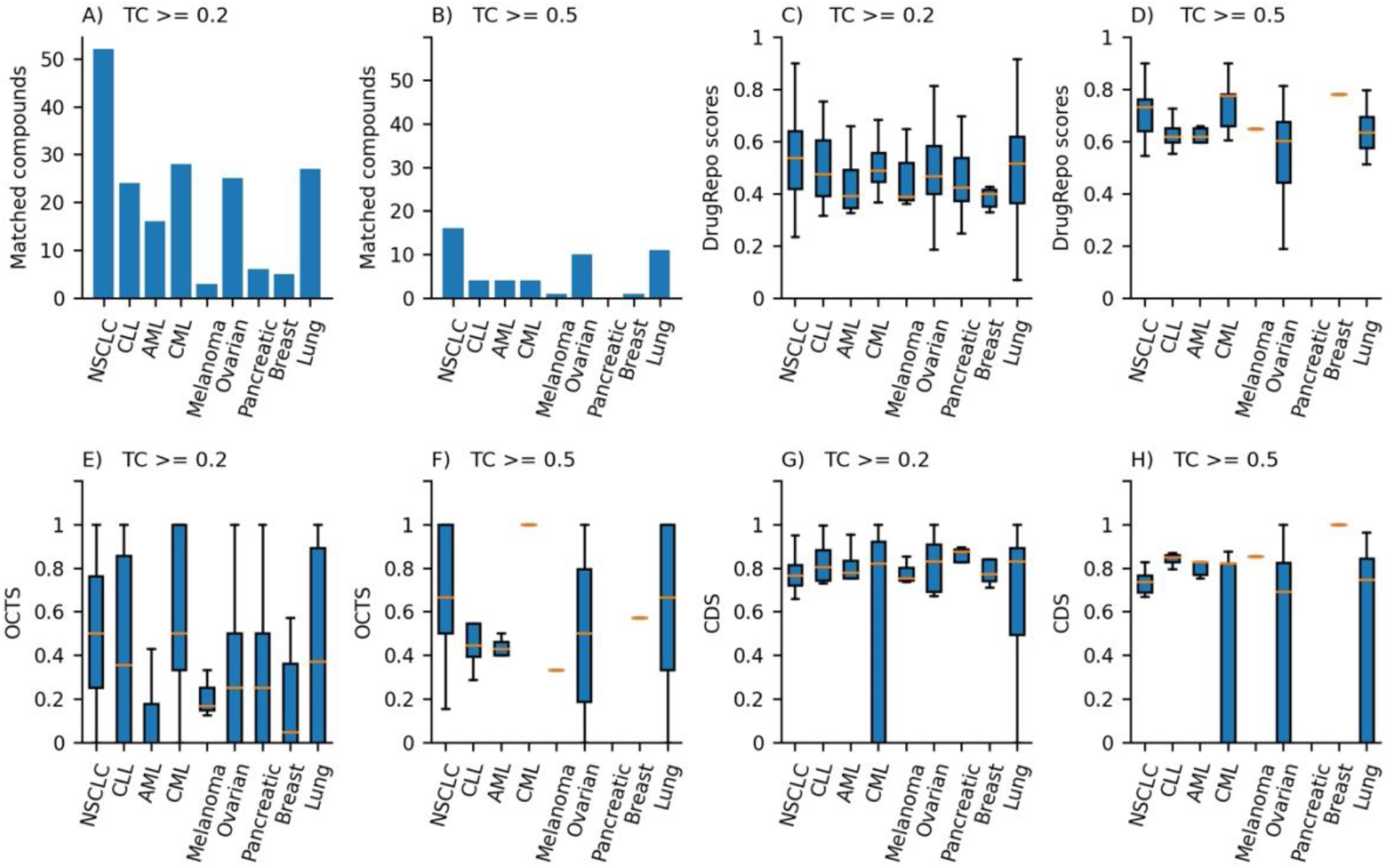
(A) The number of matched candidate compounds across nine cancer types using structural similarity (TC) >=0.2, (B) Number of matched candidate compounds using structural similarity (TC) >=0.5, (C) Distribution of DrugRepo scores for matched compounds for structural similarity (TC) >=0.2, (D) Distribution of DrugRepo scores for matched compounds for structural similarity (TC) >=0.5. (E) Distribution of compound-disease score for repurposed compounds TC >=0.2, (F) Distribution of compound-disease score for repurposed compounds TC >=0.5, (G) Distribution overlapping target profile scores between approved and repurposed compounds TC >=0.2, (H) Distribution overlapping target profile scores between approved and repurposed compounds TC >=0.5.

All these matched indications have either successfully passed phase I or phase II. There could be some candidate repurposing compounds having high DrugRepo scores but unable to match with compounds at clinical trials. This could be either due to the lack of mapping for some compounds (at clinical trials) or because only limited compounds are being tested at clinical trials. However, by looking at the matched candidate compounds, we may find a suitable threshold on the DrugRepo score to identify repurposing compounds.

Lowering the structural similarity threshold increased the chances of matching more repurposed compounds. **Figures 5A** and **5B** shows the number of matched compounds across the nine cancer types for TC >= 0.2 and TC >= 0.5, respectively. **Figures 5C** and **5D** shows the distribution of DrugRepo scores using TC >= 0.2 and TC >= 0.5. Most of the matched candidate compounds have a median DrugRepo score higher than 0.4. Hence, we suggest the users use at least DrugRepo >= 0.4 to get maximum repurposing compounds.

Furthermore, we explored the effect of structural similarity on OCTS and CDS. **Figures 5E** and **5F** show that structural similarity slightly affects the OCTS thresholds but has no significant impact on compound-disease scores (CDS). However, TC values affect the final DrugRepo score (higher structural similarity corresponds to higher DrugRepo scores), as shown in **Figures 5C** and **5D**. Matched compounds for most cancer types have median CDS >=0.7, showing the importance of CDS in defining DrugRepo (**Figure 5E** and **5F**). The median of OCTS for different cancer types is lower (0.1-0.5) than CDS because complete target profiles (across the entire druggable genome) for most of the compounds are not experimentally tested. The average number of targets for the candidate and approved compounds for each of the five databases is less than 7 ^[27]^. However, with the availability of additional high throughput DTI studies, the distribution of OCTS in DrugRepo score may also increase.

Setting a lower threshold on DrugRepo scores may result in more false positives (compounds not in clinical trials). So, we also analysed the proportion of hits across ten thresholds on DrugRepo scores. As shown in **Figure 6**, the increase in DrugRepo threshold reduces the risk of getting false positives. Especially hit percentage >=5% (for all cancers) at DrugRepo score >=0.4. However, having miss-hits with clinical trials does not necessarily mean false positives, as clinical trials contain limited number of studies/compounds.

**Figure 6:**
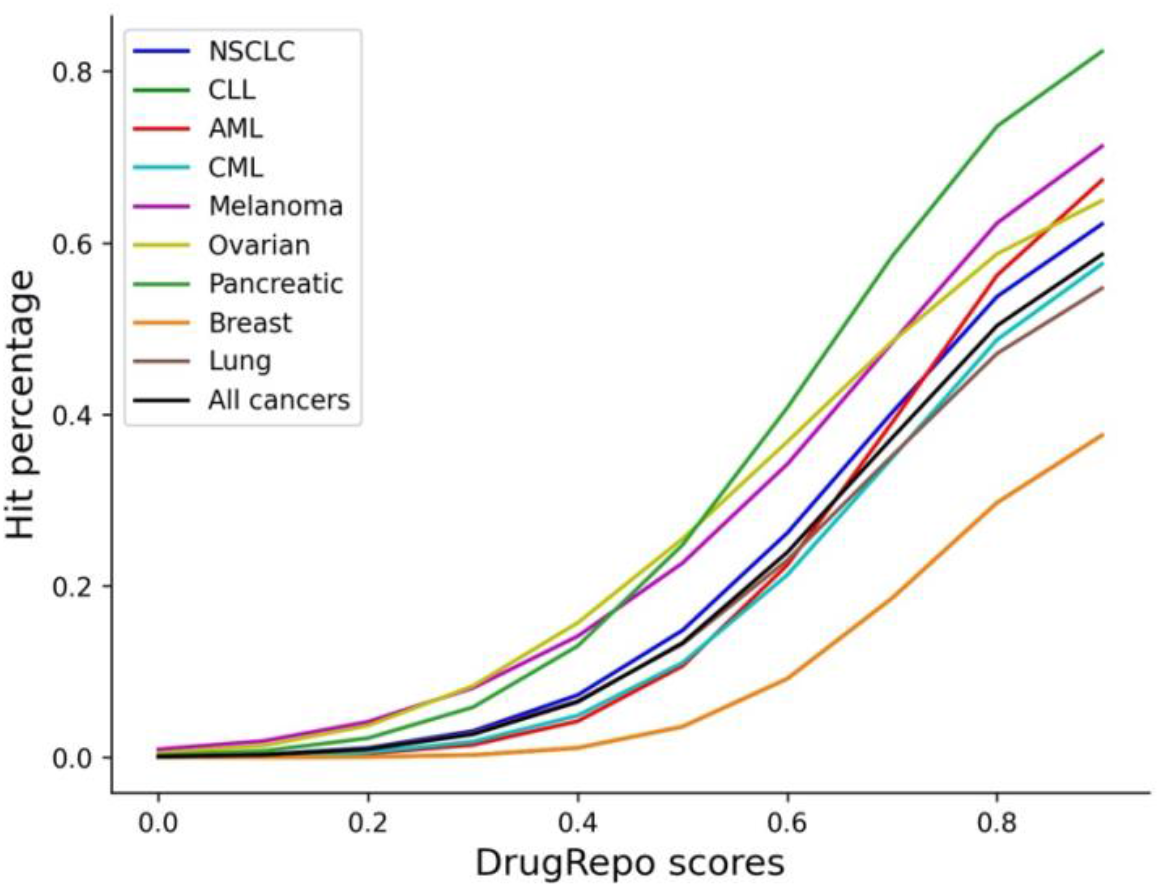
Hit ratio at different thresholds on different DrugRepo scores. The X-axis shows thresholds on DrugRepo scores, and Y-axis shows the percentage of hits while matching with repurposed compounds at https://clinicaltrials.gov/

Based on these successful matching, we can claim that a DrugRepo score >= 0.4 might guarantee the repurposing of a candidate compound with less chances of false positives. Not many compounds have been tested in clinical trials; therefore, we suggest top-scoring compounds be tested in-vitro to evaluate the significance of DrugRepo scores.

### 4.3. Using the DrugRepo’s GUI to repurpose drugs for CML

We provide a case study on Chronic Myeloid Leukaemia (CML) using the web interface at http://drugrepo.org/. DrugRepo has three approved drugs (imatinib, nilotinib, and bosutinib) for CML. Users may check one or more of these approved drugs and customize the structural similarity and DrugRepo thresholds, as shown in **Figure 7A**. The figure’s section on DrugRepo displays the results for the selected disease and associated approved compounds. Three figures can be selected from the figures dropdown list, i.e., 1) Statistics, 2) Repurposing scores, and 3) TSNE. The ‘Statistics’ displays the top targets (for approved drugs) and a list of repurposing compounds (**Figure 7B**). We used standard InChIKey identifiers for the repurposing compounds instead of names because many preclinical and investigational compounds aren’t assigned with proper names. For targets, we used UniProt IDs. The ‘Repurposing scores’ provide a 3D scatterplot with the x-axis as TC, y-axis as OCTS and z-axis as CDS scores. Each point is a pair of approved drug and repurposed compounds. Repurposing compounds are assigned with the colors similar to the most associated approved drug (**Figure 7C**). More details about diseases, approved drugs, repurposing compounds, and the three scores will show if users click on the scatter points. T-distributed stochastic neighbour embedding (TSNE) is used to visualize the 2D similarity between approved drugs (purple) and repurposing compounds (red) as shown in **Figure 7D**. The 2D similarity was computed based on ECFP4 fingerprints.

**Figure 7:**
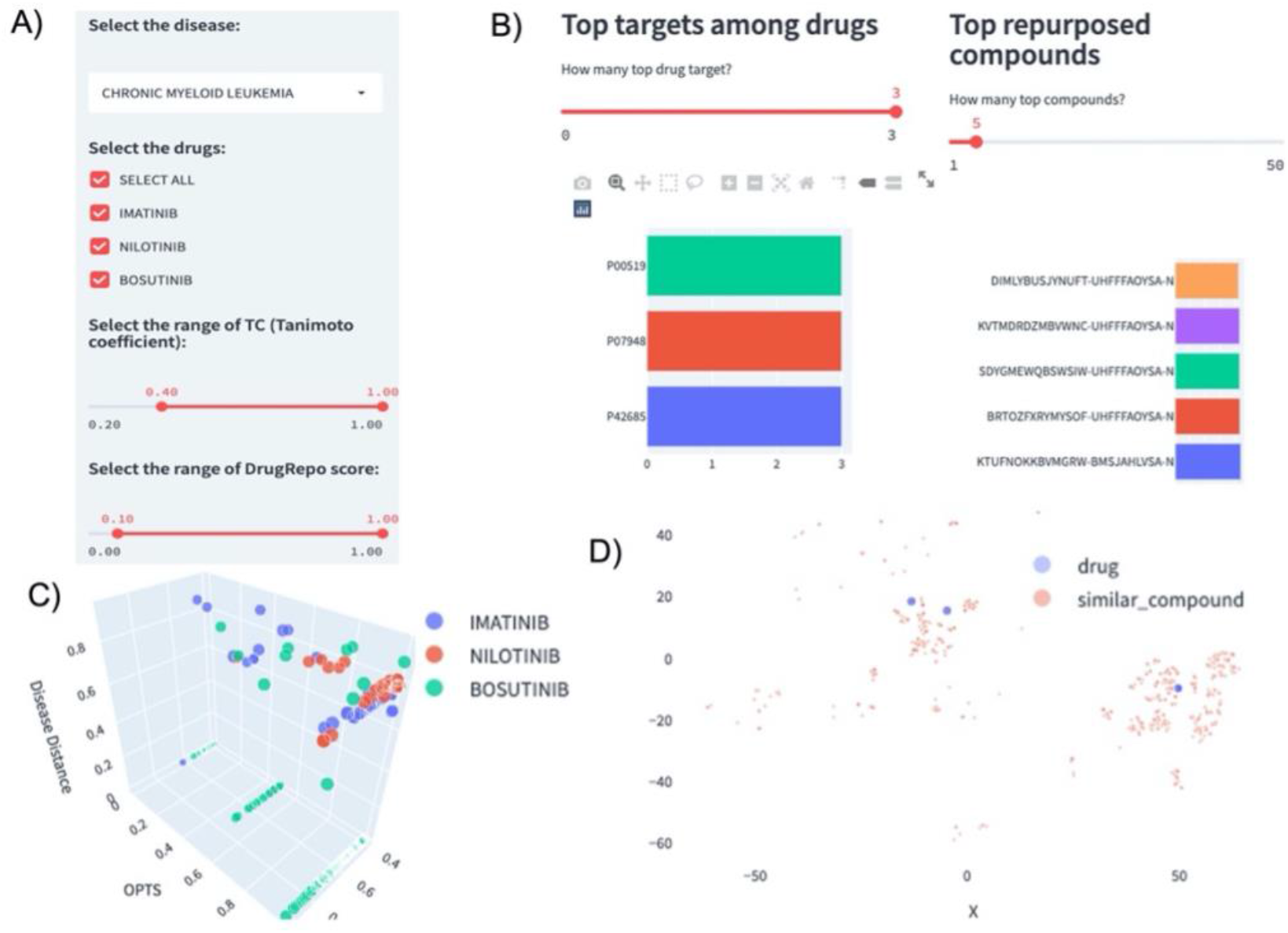
Using the DrugRepo GUI to repurpose drugs for CML. (A) Select disease interface, users can uncheck any of the approved drugs, modify TC, and DrugRepo thresholds, (B) Top targets for approved drugs and top repurposing compounds, (C) Drug repurposing plot, each point is based on three scores for an approved drug and repurposing compound, (D) The 2D visualization for structural similarities between approved drugs and repurposing compounds.

Down to the Figures, there is a Tables section containing four tables as shown in the ‘Tables’ dropdown list, 1) Drug repurposing table, 2) Approved drugs for the disease, 3) Disease-gene associations, and 4) Drug target profiles for drug and compounds. The ‘Drug Repurposing’ table provides four scores (TC, OCTS, CDS and DrugRepo) between approved and repurposing drugs. The ‘Approved drugs for the disease’ table displays UML-CUIs for the selected disease and standard InChIKeys of the approved drugs. The ‘Disease-gene associations’ table displays the genes associated with the selected disease. Finally, the ‘Drug target profiles for drug and compounds’ table provides a list of drug targets associated with the approved drugs and repurposing compounds. Users can take advantage of filter and sort options to customize the results and download data for further analysis.

## Conclusion

DrugRepo provides a platform for repurposing a massive number of compounds using chemical and genomic features. We evaluated our methods by leveraging the datasets available at clinical trials and CTD. DrugRepo is based on three components, i.e., structural similarity (TC), overlapping compound-target score (OCTS), and compound-disease scores (CDS). CDS (median ≥ 0.75 in nine cancer types) is based on PPI networks and is more effective than OCTS (median ≥ 0.3) for matching repurposed compounds (as shown in **Figures 5G & 5H**). By matching repurposed compounds, we suggest a DrugRepo score >=0.4 as the threshold to repurpose compounds for a selected disease as it has lower chances of having false positives (**Figure 6**). Computational analysis is further supported by a web application (http://drugrepo.org/), where users can select a particular disease and obtain repurposed compounds along with other parameters. Using DrugRepo GUI, users can select a particular disease, check the approved drugs, targets associated with approved drugs or candidate compounds, and finally download the compounds that can be repurposed for the selected disease. Users can adjust thresholds for structural similarity (TC) and DrugRepo scores (TC), as shown in **Figure 7A**. We also provide user-friendly visualization by which users can apply different filters to obtain specific results and finally download results for further analysis. The web application can help design new drug repurposing applications and use existing information for predictive analysis.

However, our method has some limitations. For instance, the proposed approach is dependent on OCTS between approved drugs and candidate compounds. Missing data in DTI profiles for an approved drug or candidate compound may cause failure in capturing some of the important repurposing candidates. Although we integrated DTI profiles from the five most comprehensive databases, the average number of targets for approved drugs is around seven, which is much less than expected (as druggable targets are around 1000). However, new releases of DTI databases (such as ChEMBL, DrugBank or BindingDB) may bring additional curated DTIs, resulting in better repurposing applications. We will keep updates with the newly curated datasets to make DrugRepo more effective. Presently DrugRepo is based only on three components. However, with more components (such as gene expression data), results can be further improved. We will therefore incorporate these improvements in the next version of DrugRepo.

## Key points

- We proposed a novel scoring algorithm for repurposing huge collection of pre-clinical compounds. The analysis is supported by web tool available at: http://drugrepo.org/
- DrugRepo score is based on three components i.e. molecular structural similarity (TC), Overlapping compound-target score (OCTS) and compound-disease score (CDS).
- DrugRepo GUI helps translational researchers to design new drug repurposing applications and to perform predictive analysis

## DECLARATIONS

### Availability of data and material

DrugRepo is available at http://drugrepo.org/.

### Authors’ contributions

Conceptualization: Z.T, J.T; Methodology: Z.T, Y.W; Software: Y.W, J.A, Y.H; Formal analysis: Y.W, H.Y, Z.T; Writing: J.T, Z.T, Y.W; Review & Editing: M.V; Supervision: J.T, Z.T

## Acknowledgements

We thank CSC, Finland for providing us the IT services.

**Yinyin Wang** is a PhD student at university of Helsinki. She is developing methods for network pharmacology modeling for herb medicine.

**Jehad Aldahdooh** is a PhD student at university of Helsinki. He is developing text mining applications for drug target interactions.

**Yingying Hu** is a master’s student at university of Helsinki. She is doing research on systems biology applications.

**Hongbin Yang** is a postdoc researcher at university of Cambridge. He is doing research on systems biology applications.

**Markus Vähä-Koskela** is senior researcher at Institute for Molecular Medicine Finland. He is doing research in areas such as: onco-immunology, immunotherapy, and translational cancer medicine.

**Jing Tang** is assistant professor at university of Helsinki. He is working on mathematical, statistical and informatics tools to tackle biomedical questions.

**Ziaurrehman Tanoli** is senior researcher at university of Helsinki. His research is mostly focused on computational drug repurposing. He is also developing bioinformatics tools for drug target interactions.

